# Polθ is phosphorylated by Polo-like kinase 1 (PLK1) to enable repair of DNA double strand breaks in mitosis

**DOI:** 10.1101/2023.03.17.533134

**Authors:** Camille Gelot, Marton Tibor Kovacs, Simona Miron, Emilie Mylne, Rania Ghouil, Tatiana Popova, Florent Dingli, Damarys Loew, Josée Guirouilh-Barbat, Elaine Del Nery, Sophie Zinn-Justin, Raphael Ceccaldi

## Abstract

DNA double strand breaks (DSBs) are deleterious lesions that challenge genome integrity. To mitigate this threat, human cells rely on the activity of multiple DNA repair machineries that are tightly regulated throughout the cell cycle^1^. In interphase, DSBs are mainly repaired by non-homologous end joining (NHEJ) and homologous recombination (HR)^2^. However, these pathways are completely inhibited in mitosis^3–5^, leaving the fate of mitotic DSBs unknown. Here we show that DNA polymerase theta (Polθ)^6^ repairs mitotic DSBs and thereby maintains genome integrity. In contrast to other DSB repair factors, Polθ function is activated in mitosis upon phosphorylation by the Polo-like kinase 1 (PLK1). Phosphorylated Polθ is recruited to mitotic DSBs, where it mediates joining of broken DNA ends, while halting mitotic progression. The lack of Polθ leads to a shortening of mitotic duration and defective repair of mitotic DSBs, resulting in a loss of genome integrity. In addition, we identify mitotic Polθ repair as the underlying cause of the synthetic lethality between Polθ and HR. Our findings reveal the critical importance of mitotic DSB repair for maintaining genome stability.

## Introduction

Cells can enter mitosis with DSBs that are formed during interphase^7^, and DSBs can also form in mitosis, as a consequence of replication stress (RS)^8,9^. As a result, HR-deficient (HRD) cells, that experience elevated levels of RS, are prone to accumulate mitotic DSBs^10^. It has been shown that broken DSB ends can be held together by a tethering complex and pass through mitosis in this manner, awaiting repair in the next cell cycle^11,12^. However, this pathway alone might not be sufficient in safeguarding the genome against mitotic DSBs, arguing for the existence of parallel alternatives. A heretofore unaddressed, yet intriguing possibility is that DSB repair pathways, alternative to HR and NHEJ^13,14^, might be active in mitosis. The DNA polymerase theta (Polθ) has recently emerged as a major player in alternative end joining (alt-EJ) repair^15–17^, essential for HR-deficient cell survival^18–21^. Here we reasoned that Polθ-mediated alt-EJ could play a key role in repairing mitotic DSBs, thereby maintaining genome integrity and ensuring HR-deficient cell survival.

## Results

### Polθ functions downstream of the HR pathway in S phase

To investigate the function of Polθ in human cells, we tagged the endogenous POLQ locus with a NeonGreen (NG) tag, or exogenously expressed GFP-tagged Polθ in several human cell line models. In these cells, Polθ localized to sites of DSBs in a poly (ADP-ribose) polymerase (PARP)-dependent manner, as previously reported^18,19^ (Extended Data Fig. 1a-d). In order to decipher the regulation of Polθ localization to DSBs, we performed an unbiased immunofluorescence screening of Polθ foci formation. Briefly, first, RPE-1 cells expressing GFP-Polθ were transfected with an siRNA library targeting factors involved in the DNA damage response (DDR). Cells were exposed to ionizing radiation (IR) and Polθ foci were scored by automatic fluorescence microscopy. As expected, siRNAs against Polθ itself or FANCD2^22^ abolished Polθ foci formation (Fig. 1a). Surprisingly, knockdown of core factors in HR, such as BARD1, BRCA1, PALB2 or BRCA2 resulted in a striking reduction of Polθ foci formation (Fig. 1a and Table 1). Results of the screen were validated by independent, siRNA or mAID (degron)-mediated knockdown of various HR genes (Extended Data Fig. 1e,f). Additionally, immunoprecipitation (IP) coupled to label-free mass spectrometry (MS) analysis of tagged Polθ showed that Polθ co-purified with several members of the HR pathway, such as BARD1, BRCA1, PALB2, and BRCA2 (Extended Data Fig. 1g-i and Table 2). Altogether, our data provides evidence that Polθ is recruited to sites of DSBs downstream of the HR pathway in interphase. These results are in accordance with recent studies demonstrating that Polθ and HR effectors, such as RAD51 and RPA, can compete for the repair of similar substrates^18,23,24^ (Extended Data Fig. 1j).

**Figure 1.**
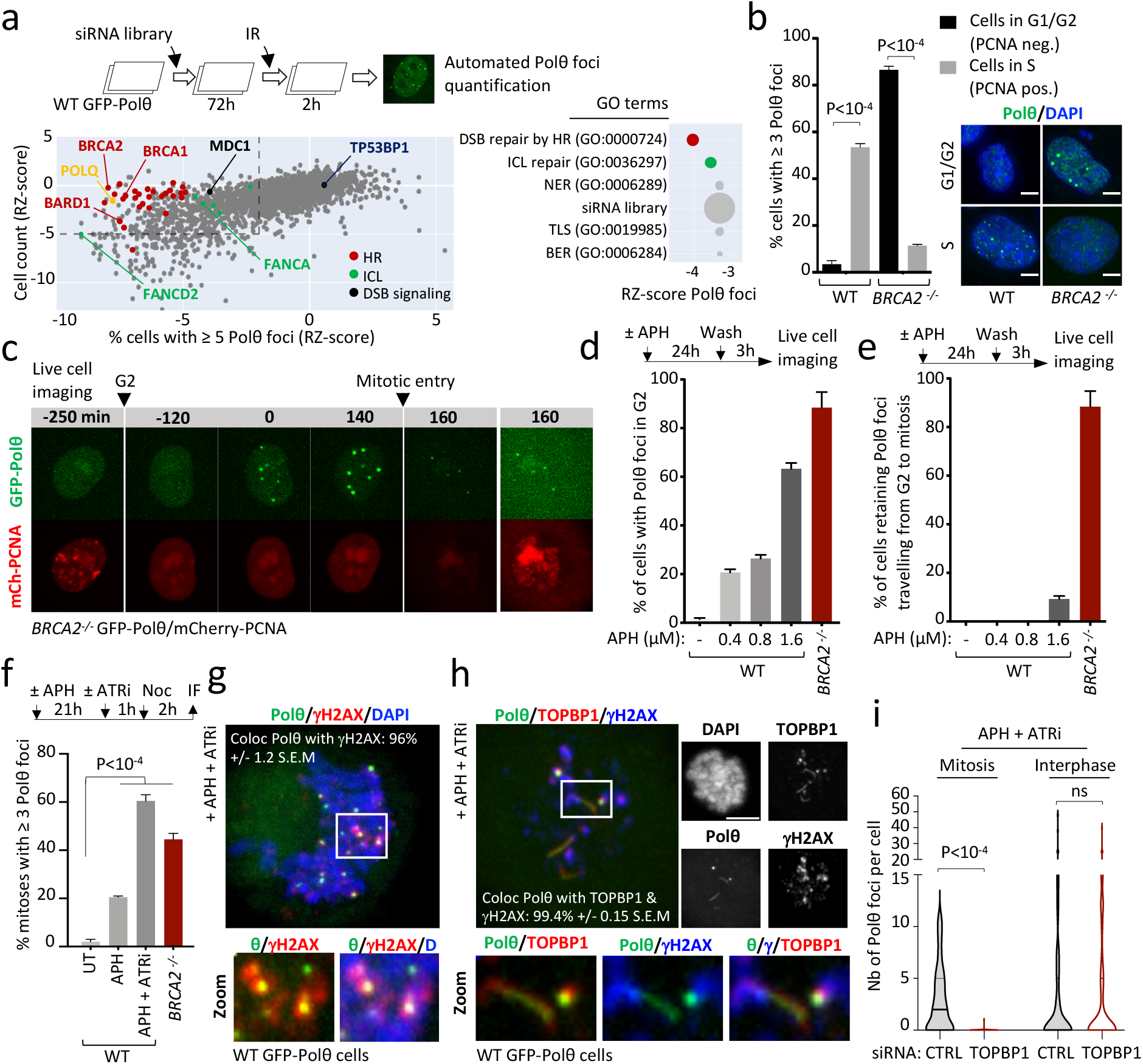
Polθ has HR-dependent and -independent functions in different cell cycle phases. **a**, siRNA screen for IR-induced Polθ foci formation. Schematic of the screen is shown on top. siRNAs against indicated targets were plotted as Robust Z-score (RZ-score) according to cell survival (y axis) and Polθ foci formation (x axis). For each siRNA, median RZ-score of each of the three replicate experiments is shown. Arrows show the strongest RZ-score of indicated gene. Enriched GO terms of biological processes, identified among all targets are shown on the right. HR = Homologous Recombination, ICL = Interstrand Crosslink repair, NER = Nucleotide Excision Repair, TLS = Translesion Synthesis and BER = Base Excision Repair. **b**, Cell cycle distribution of Polθ foci in wild-type (WT) and *BRCA2^-/-^* cells. **c**, Representative images of live microscopy analysis of *BRCA2^-/-^* cells expressing GFP-Polθ and mCherry-PCNA. (n > 200 cells per condition). **d**, Quantification of Polθ foci in G2 in WT and *BRCA2^-/-^* cells upon indicated doses of aphidicolin (APH). Schematic of the experiment is shown on top. (n > 75 cells per condition). **e**, Quantification of cells retaining Polθ foci while travelling from G2 to mitosis. **f**, Quantification of Polθ foci in mitotic cells upon indicated treatment (APH: 0.4 μM, 24 h). **g**, **h**, Representative images, and quantification of Polθ foci (g) and filaments (h) colocalization with γH2AX and TOPBP1 in mitosis. **i**, Quantification of Polθ foci formation upon indicated treatment in interphase or mitotic cells. (n > 50 mitotic cells and > 100 interphase cells per condition). Scale bars represent 5 μm. Data represent at least three biological replicates. Data show mean +/- S.E.M., except violin plots (i) showing median with quartiles. Asterisks indicate statistically significant values (*P <0.05; **P <10^-2^, ***P <10^-3^).

### Polθ forms foci at mitotic entry, independently of HR

To decipher the conundrum of HR-dependent Polθ foci formation and synthetic lethality between Polθ and HR^18,19^, we compared Polθ foci formation in cells lacking HR with wild-type (WT) isogenic counterparts^20^. We found that, the cell cycle distribution of Polθ foci in *BRCA2^-/-^* cells was the inverse of that of WT cells. In WT cells, Polθ foci were mostly visible in S phase (PCNA-positive) cells, while in *BRCA2^-/-^* cells, Polθ foci were prominent in G1 and G2 (PCNA-negative) cells but absent in S (Fig. 1b and Extended Data Fig. 1k and 2a, Videos 1-3). Our data suggests that, while Polθ functions downstream of HR in S, it performs HR-independent functions from G2 to the next G1 phase. Using live-microscopy, we found that, in *BRCA2^-/-^* cells, Polθ accumulated in G2, close to mitotic entry (after disappearance of PCNA foci) (Fig. 1c). Additionally, Polθ foci in G2 could be induced by replication stress (RS) (aphidicolin (APH) treatment) in WT cells, to the level of *BRCA2^-/-^* cells (Fig. 1d). Interestingly, the vast majority of Polθ foci in *BRCA2^-/-^* cells were transmitted from G2 to mitosis (Fig. 1e). Taken together, our findings indicate that, in *BRCA2^-/-^* cells, Polθ marks intrinsic RS-induced lesions in G2 that remain unresolved and are transmitted to mitosis.

### Polθ is recruited to mitotic DSBs

We found that, similar to G2 cells, mitotic cells showed an increase in Polθ foci formation upon RS, confirming the persistence of Polθ-marked lesions (Fig. 1f and Extended Data Fig. 2b-d). Almost all mitotic Polθ foci colocalized with phosphorylated histone H2AX (γH2AX, a DSB marker), indicating that Polθ marks RS-induced DSBs in mitosis (Fig. 1g and Extended Data Fig. 2c,d). Of note, Polθ colocalized poorly with mitotic DNA synthesis (MiDas) foci (labelled by EdU and FANCD2)^25^ arguing for a MiDas-independent role of Polθ (Extended Data Fig. 2e,f). While HR and NHEJ repair are inactivated in mitosis, early events of the DDR still occur, such as recruitment of the scaffold proteins MDC1 and TOPBP1 to DSBs, in an ATM-dependent manner^11,26,27^. We found that, the vast majority of mitotic Polθ foci co-localized with MDC1 and TOPBP1 foci, while Polθ foci in interphase co-localized only with MDC1 but not with TOPBP1 (Extended Data Fig. 2g). Interestingly, Polθ also formed filament-like structures perfectly associating with TOPBP1 (Fig. 1h and Extended Data Fig. 2h,i). This was particularly evident in cells under RS, suggesting that these mitotic filaments are instrumental to the cellular response to RS (Extended Data Fig. 2h,i). Furthermore, MDC1 knockdown prevented the formation of Polθ foci in both interphase and mitosis, while TOPBP1 knockdown suppressed Polθ foci (and filament) formation only in mitosis (Fig 1i and Extended Data Fig. 2j). This shows that, while MDC1 acts similarly in both interphase and mitotic DDR, the interaction between TOPBP1 and Polθ is specific to mitosis.

To further confirm the recruitment of Polθ to mitotic DSBs, we irradiated mitotic cells (collected by gentle shake-off of nocodazole-arrested cells) and measured Polθ foci formation. We found that Polθ foci formation could be induced upon IR (Extended Data Fig. 3a-d). This was especially striking, since there are no other known DNA repair factors that can be recruited to DSBs generated in mitosis^1^. Furthermore, IR-induced mitotic Polθ foci (similarly to RS-induced) co-localized with the TOPBP1/MDC1 complex and relied on ATM and MDC1 for their formation (Extended Data Fig. 3e-i). Taken together, our data suggests that the early steps of the DDR mediated by ATM and MDC1 is conserved between interphase and mitosis. However, repair pathways diverge downstream of MDC1, with classical DSB repair (HR and NHEJ) dominating in interphase and, TOPBP1/Polθ comprising a novel, alternative pathway responding to mitotic DSBs.

### Polθ repairs mitotic DSBs

In mitosis, while it is well accepted that HR and NHEJ repair activities are inhibited, the activity of alternative pathways remains unaddressed. While there has not been direct evidence of DSB repair in mitosis, recent data suggests that alt-EJ^28,29^ and Polθ could be active in mitosis^30–32^. To detect DSB repair in mitosis, we took advantage of recent advances in genome editing, allowing the induction of DSBs within minutes after Cas9 protein transfection. Nocodazole-arrested cells were transfected with a Cas9/gRNA complex to induce DSB formation at two different sites in the AAVS1 safe locus, whose mutagenic repair would destroy a nearby HphI restriction site. After DSB induction, a small genomic region around the cut site was PCR amplified and digested by HphI. Finally, undigested PCR products (representing mutagenic repair) were subjected to Sanger sequencing (Fig. 2a and Extended Data Fig. 4a).

**Figure 2.**
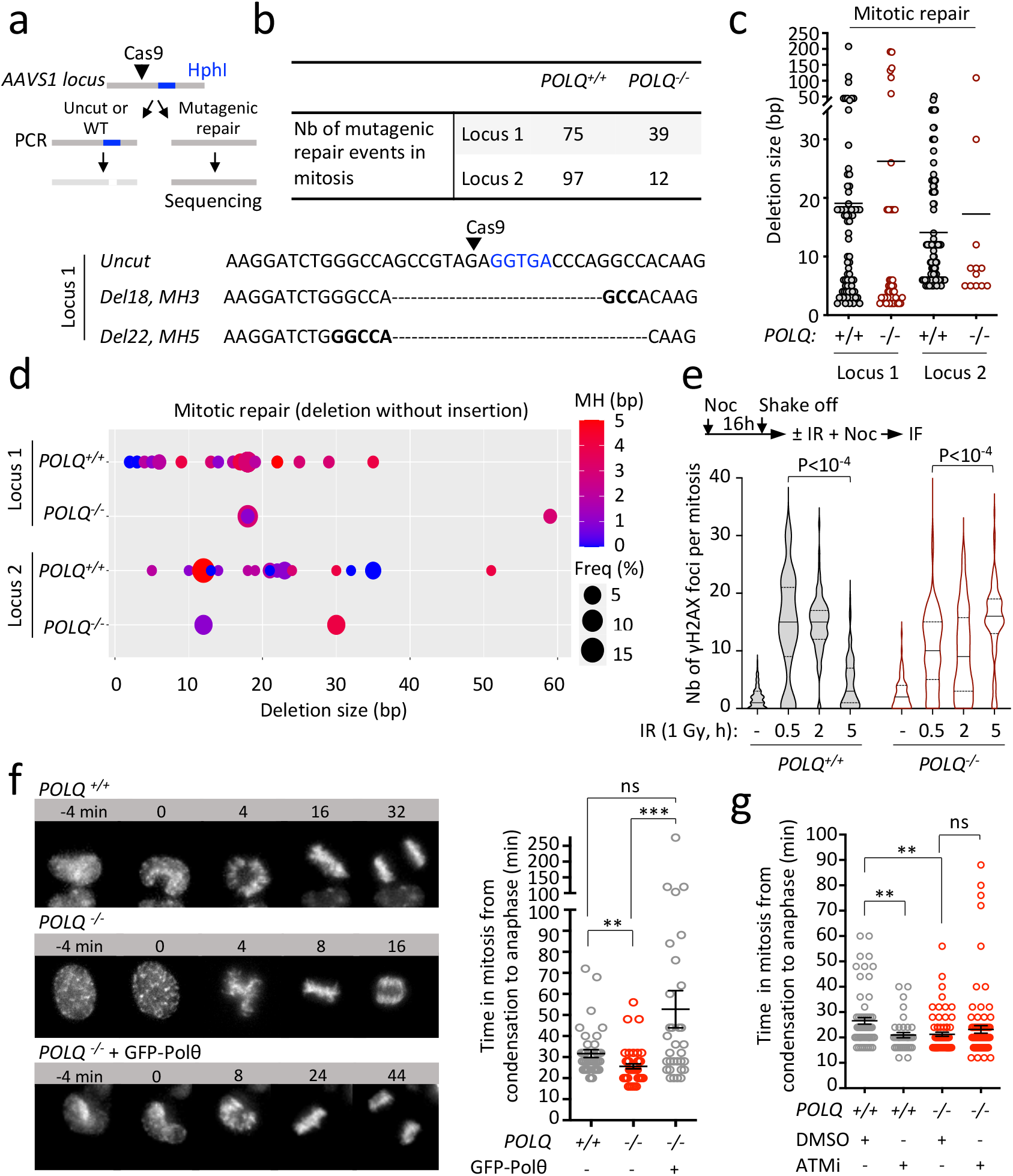
Polθ repairs DSBs in mitosis and controls mitotic timing. **a,** Experimental workflow. **b**, Table recording the total number of mitotic DNA repair events identified following CRISPR-Cas9 induced-cleavage in indicated cell lines. Representative repaired DNA sequences. Deletions (Del) and microhomologies (MH) size are indicated. At repair junction, MH are indicated in bold and Hph1 recognition site in blue letters. **c**, Deletion size of mitotic DNA repair events identified in WT and *POLQ^-/-^* cells. Each dot represents a mitotic DNA repair event. **d**, Graph showing the frequency, deletion size and microhomology (MHS) use of mitotic DNA repair events identified in WT and *POLQ*^-/-^ cells. Events with deletions size < 60 bp are represented. **e,** Quantification of γH2AX foci at different time points after radiation (1 Gy) in mitosis. (n > 80 mitotic cells per condition). **f**, **g**, Representative images and quantification of mitotic duration in indicated cell lines labeled with SiR-DNA and treated with the ATM inhibitor (ATMi) when indicated. (n > 40 mitotic cells per condition). Scale bars represent 5 μm. Data represent at least three biological replicates. Data show mean +/- S.E.M., except violin plots (e) showing median with quartiles. Asterisks indicate statistically significant values (*P <0.05; **P <10^-2^, ***P <10^-3^).

We observed evidence of mutagenic mitotic DSB repair at both of the two loci tested (Fig. 2b). To rule out contamination from interphase cells, the purity of mitotic samples was confirmed by histone H3 phospho-S10 immunofluorescence (H3pS10) (a specific marker of mitosis) (96-100% purity)(Extended Data Fig. 4b). We next compared mitotic DSB repair to that of asynchronous control cells. We found that repair products obtained from cells in mitosis differed significantly from those in asynchronous cells (more products with deletions > 10 bp associated with the use of microhomology (MH)), further ruling out the hypothesis of a contaminant (Extended Data Fig. 4c-e).

Furthermore, we observed a striking difference in mitotic DSB repair between WT and *POLQ knockout* (*POLQ^-/-^*) cells (Fig. 2b-d and Extended Data Fig. 4f-g, Table 3). The number of repair events was greatly diminished at both loci in *POLQ^-/-^* cells, indicating that Polθ is required for efficient mitotic DSB repair (Fig. 2b). Previous studies have demonstrated that Polθ-mediated repair uses microhomologies (MHs) and generates deletions of up to 60 bp^33–36^. Accordingly, we found that, while WT and *POLQ^-/-^* cells both exhibited repair products with <10 or >50 bp deletions, repair products with deletions ranging from 10 to 50 bp were remarkably absent in *POLQ^-/-^* cells (Fig 2c, d). In addition, we showed that, the use of MH was common in WT cells but greatly diminished in the few repair products detected in *POLQ^-/-^* cells (Extended Data Fig. 4f). Finally, to assess the contribution of Polθ to mitotic DSB repair in a more quantitative manner, we determined the kinetics of γH2AX foci resolution after IR in mitosis. We found that, while the vast majority of γH2AX foci in WT cells were resolved 5 hours (h) after IR, they persisted in *POLQ^-/-^* cells (Fig. 2e). Altogether, our results pinpoint Polθ as a major player in mitotic DSB repair. As opposed to it being an alternative repair factor in interphase, we posit that Polθ is the main DSB repair factor in mitosis.

### Polθ controls mitotic timing

In interphase, DNA damage is accompanied by the activation of checkpoints, halting cell cycle progression, thus allowing time for DNA repair. It has been shown that DSBs in mitosis delay anaphase onset and that ATM and MDC1 can participate in this phenomenon, arguing for the existence of a DNA damage-induced checkpoint in mitosis^37,38^. Since Polθ is recruited to mitotic DSBs in an ATM- and MDC1-dependent manner, we wondered whether Polθ could be a part of this checkpoint.

First, we found that, Polθ is recruited to DSBs localized at regions known to trigger checkpoint activation in mitosis (centromeric regions (CREST-positive) and acentric DNA fragments)^38,39^ (Extended Data Fig. 5a-d). Second, we measured time spent in mitosis as a proxy for mitotic checkpoint activation. Time lapse microscopy revealed that *POLQ^-/-^* cells spend less time in mitosis than WT counterparts, which can be rescued by complementation with WT Polθ. Altogether, this points towards a role of Polθ in regulating mitotic duration (Fig. 2f and Extended Data Fig. 5e, Videos 4-6). In addition, we found that while ATMi induced a shortening of mitotic duration, as previously reported^38^, it did not further impact mitotic duration in *POLQ^-/-^* cells indicating epistasis between Polθ and ATM in mitotic checkpoint activation (Fig. 2g). Altogether, our data support a role for Polθ in sensing mitotic DSBs to slow down cell cycle progression and allow DNA repair.

### Polθ is phosphorylated by PLK1 in mitosis

We next sought to elucidate the regulation of mitotic Polθ activity. The mitotic kinases CDK1 and PLK1^40^ restrict DSB repair through the phosphorylation of several NHEJ and HR factors, such as 53BP1 and BRCA2^3,4,41^. A 53BP1 mutant (T1609A, S1618A), which cannot be phosphorylated by PLK1, escapes negative regulation and forms IR-induced foci in mitosis^4,39^. Interestingly, we found that, this unrestrained 53BP1 foci formation abolished Polθ foci formation in mitosis (Extended Data Fig. 6a). This suggests a competition between the two pathways and PLK1 as a mediator of pathway choice. We also observed a colocalization between Polθ and PLK1 foci (Extended Data Fig. 6b). Together, these prompted us to test a potential phosphorylation of Polθ by PLK1 in mitosis.

To investigate this, we immunoprecipitated (IP) Polθ and assessed phosphorylation by immunoblot analysis (using pan phospho antibodies). We observed a phosphorylation signal corresponding to the size of Polθ when IP was performed from mitotic cell extracts. In contrast, this signal was completely absent from asynchronous cells, suggesting that Polθ is only phosphorylated in mitosis (Fig. 3a and Extended Data Fig. 6c,d). Importantly, this phosphorylation signal was abolished when cells were treated with two different PLK1 inhibitors (PLK1i), indicating that PLK1 is responsible for Polθ phosphorylation in mitosis (Fig. 3a and Extended Data Fig. 6d). Furthermore, *in vitro* incubation of Polθ-immunoprecipitations with purified recombinant PLK1 enzyme in the presence of radioactive ATP confirmed a direct phosphorylation of Polθ by PLK1 (Extended Data Fig. 6e).

**Figure 3.**
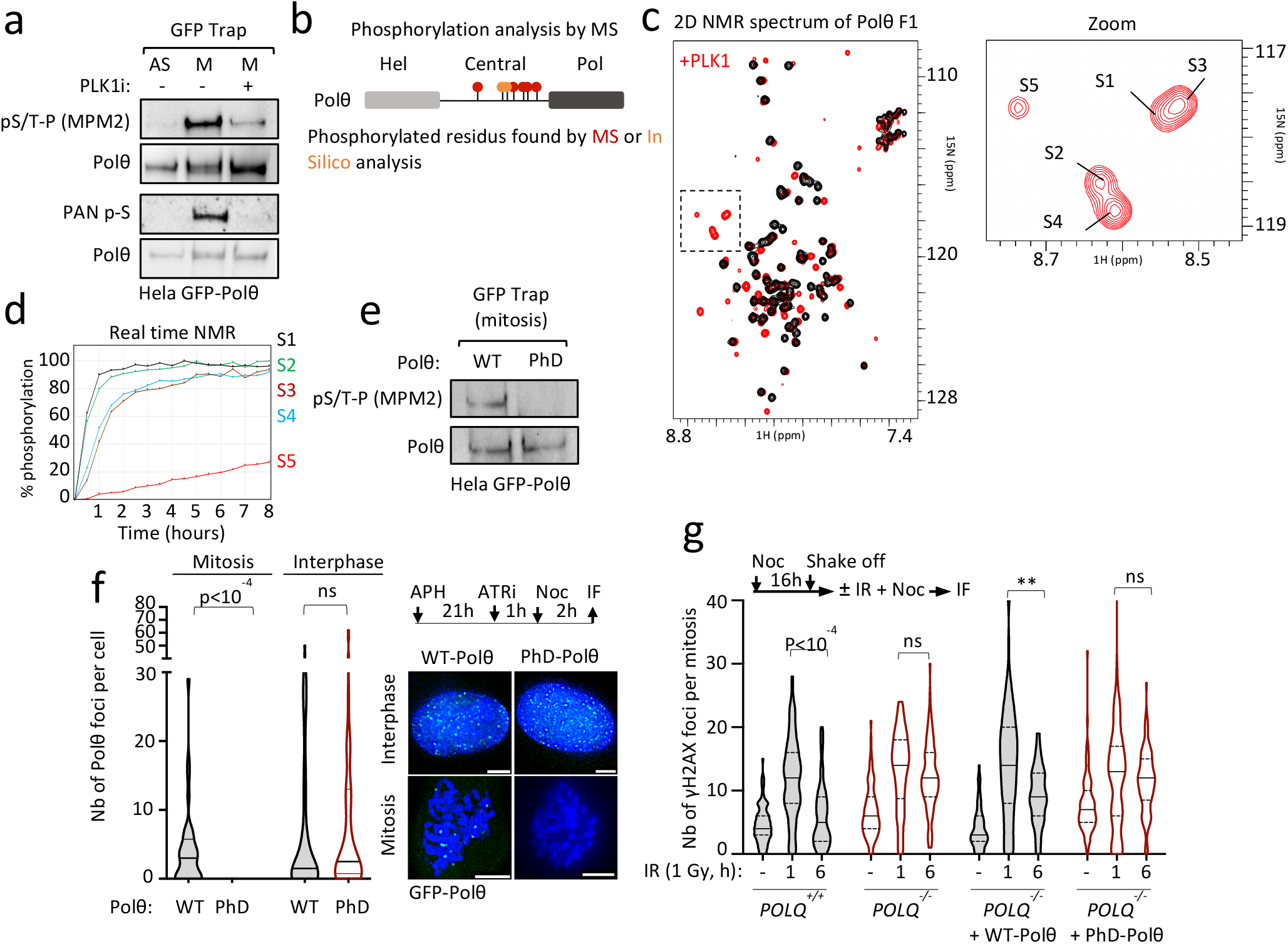
Polθ is phosphorylated by PLK1 in mitosis. **a**, Immunoblot analyses following immunoprecipitations of Polθ-GFP from asynchronous (AS) and mitotic cells. **b**, Scheme depicting PLK1 phosphorylation sites on Polθ identified by quantitative mass spectrometry (red) and *in silico* analysis (orange). **c**, Superposition of the 2D NMR ^1^H-^15^N SO-FAST HMQC spectra recorded on an ^15^N, ^13^C labeled Polθ fragment (Polθ F1), before (black) and after (red) incubation with PLK1. A zoom shows the spectral region containing the NMR signals of phosphorylated residues. **d**, Phosphorylation kinetics, as monitored by real-time NMR. Intensities of the NMR peaks corresponding to phosphorylated residues shown in (c) were measured on a series of SO-FAST HMQC spectra recorded on the Polθ fragment incubated with PLK1 and plotted as a function of time. **e**, Immunoblot analyses following immunoprecipitation of WT- or PhD-Polθ from mitotic cells. **f**, Representative images and quantification of Polθ foci in cells expressing WT- or PhD-Polθ. **g**, Quantification of γH2AX foci at indicated time points after radiation (1 Gy) in mitosis in indicated cell lines. Scale bars represent 5 μm. Data (f, g) represent at least three biological replicates. Data show mean +/- S.E.M., except violin plots (g, i) showing median with quartiles. Asterisks indicate statistically significant values (*P <0.05; **P <10^-2^, ***P <10^-3^).

In order to identify the Polθ residues phosphorylated by PLK1, we performed quantitative mass spectrometry (MS)-based phosphorylation analysis of Polθ in mitosis with or without PLK1i. This analysis found 5 sites phosphorylated by PLK1, including a predicted PLK1 phosphorylation site^42,43^ (Fig. 3b and Table 4). We next applied nuclear magnetic resonance (NMR) spectroscopy to a recombinant peptide (Polθ F1) containing two predicted phosphorylation sites undetectable by MS. NMR analysis of the Polθ fragment purified from *E. coli* identified 5 serines as phosphorylated by PLK1 *in vitro* (Fig. 3c and Extended Data Fig. 6f). The fast kinetics of Polθ phosphorylation observed by NMR suggests that Polθ is a preferential substrate for PLK1 (Fig. 3d). All phosphorylation sites were conserved in evolution, even though they are located within the large, central, non-conserved region of Polθ. This is particularly striking for 4 phosphorylation sites, that are clustered in a region highly conserved within vertebrates, thus indicating a crucial function for these residues (Extended Data Fig. 6g and Table 5).

### Polθ phosphorylation by PLK1 controls mitotic DSB repair

Next, to assess the functional consequences of Polθ phosphorylation by PLK1, we mutated all serines that are conserved canonical phosphorylation sites, and serines identified by MS and NMR, to alanines and complemented *POLQ^-/-^* cells with either WT or non-phosphorylable (PhD) Polθ (Fig. 3e and Table 5). We found that while PhD-Polθ shows no defect in interphase foci formation, it fails be recruited to RS-induced DSBs in mitosis (Fig. 3f and Extended Data Fig. 7a, Video 7). Similarly, PLK1 inhibition prevented Polθ foci formation in mitosis but not in interphase (Extended Data Fig. 7b,c). In addition, Polθ filament structures were abolished in PhD-Polθ expressing cells (Extended Fig. 7d). To evaluate the role of Polθ phosphorylation on mitotic DSB repair, we determined the kinetics of γH2AX foci resolution after IR in mitosis in *POLQ^-/-^* cells complemented with either WT- or PhD-Polθ. Importantly, expression of WT-Polθ but not PhD-Polθ rescued the accumulation of mitotic DSBs observed in *POLQ^-/-^* cells (Fig. 3g). Altogether, our data shows that PLK1 acts as a positive regulator of mitotic DSB repair by Polθ, contrary to its inhibitory role of HR and NHEJ.

TOPBP1 has 9 BRCT repeats, and several of these repeats interact with phosphorylated peptides^44^. We have shown that, TOPBP1 and Polθ colocalize perfectly in mitosis, and that, TOPBP1 controls Polθ recruitment to mitotic DSBs. Furthermore, all identified phosphorylation sites on Polθ are within a region that is disordered (as predicted by AlphaFold^45^), and thus accessible for mediating protein-protein interactions (Extended Data Fig. 7e). Based on these observations, we speculate that TOPBP1 recruits Polθ in mitosis, by binding, through its BRCT domains, to phosphorylated residues of Polθ.

### Polθ forms nuclear bodies (NBs) in G1

The persistence of unreplicated DNA can result in the formation of 53BP1 nuclear bodies (53BP1 NBs) and micronuclei (MN) in the next G1 phase^46^. 53BP1 NBs represent a second opportunity to repair these lesions, in case MiDas proves insufficient. We found that, similarly to 53BP1, Polθ formed large, replication stress-induced nuclear bodies (Polθ NBs) and localized to some MN in G1 (Fig. 4a,b and Extended Data Fig. 8a-c). Polθ NBs were frequently associated with γH2AX, but poorly co-localized with 53BP1 (Fig. 4a and Extended Data Fig. 8a,b). By following each Polθ NB from formation to resolution by time lapse microcopy, we found that, in contrast to 53BP1 NBs which are resolved in S phase^47^, Polθ NBs were resolved in G1 (Extended Data Fig. 8d and Video 8). In addition, *POLQ^-/-^* cells exhibited elevated levels of 53BP1 NBs, and, conversely, 53BP1 knockdown increased Polθ NB formation (Extended Data Fig. 8e-h). Taken together, our data suggests that mitotic Polθ repair and MiDas act as parallel pathways to mitigate replication stress and repair unresolved lesions in mitosis or during the next G1.

**Figure 4.**
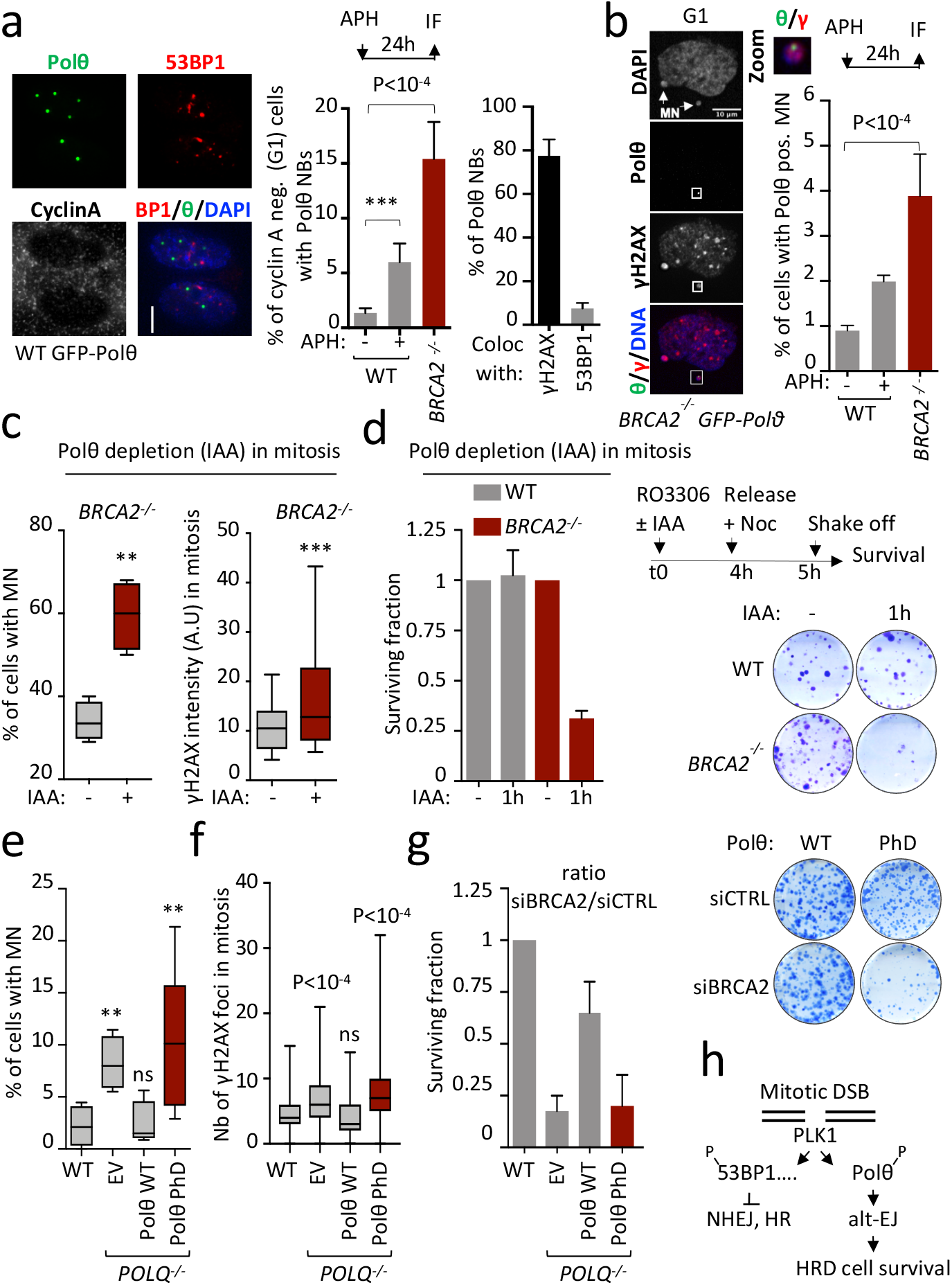
Polθ function in mitosis maintains genome stability and HR-deficient cell survival. **a**, Representative images and quantification of Polθ nuclear bodies (Polθ NBs) in cyclin A-negative (G1) cells. Colocalization of Polθ NBs with indicated proteins. **b**, Representative images and quantification of Polθ positive micronuclei (MN). **c**, Quantification of micronuclei (MN) and γH2AX intensity upon Polθ depletion in mitosis. Polθ depletion is achieved by indole-3-acetic acid (IAA) treatment (degron). **d**, Experimental workflow, representative images and quantification of 14-days clonogenic survival assays upon depletion of Polθ (IAA treatment) in mitosis. **e**, **f**, **g**, Quantification of micronuclei (MN) (e), γH2AX foci (f), and cell survival in *POLQ^-/-^* cells complemented with WT- or PhD-Polθ. **h**, Model for function of Polθ in mitosis. Scale bars represent 5 μm. Data represent at least three biological replicates. Data show mean +/- S.E.M., except box plots (c, e, f) showing median with minimum and maximum values. Asterisks indicate statistically significant values (*P <0.05; **P <10^-2^, ***P <10^-3^).

### Polθ phosphorylation in mitosis is required for genome integrity and HR-deficient cell survival

Finally, we evaluated the importance of Polθ function in mitosis in maintaining genome integrity and HR-deficient cell survival. To that end, we used the mAID system to deplete Polθ specifically in mitosis in WT and *BRCA2^-/-^* cells (Extended Fig. 8i). We found that mitotic Polθ depletion led to an increased number of unrepaired mitotic DSBs, as measured by γH2AX foci in mitosis and MN in the next G1 phase (Fig. 4c and Extended Fig. 8j,k). We also found that Polθ depletion in mitosis killed *BRCA2^-/-^* without affecting the survival of WT cells (Fig. 4d and Extended Fig. 9a). Taken together, our data highlights the crucial role of mitotic Polθ repair in maintaining genome integrity and HR-deficient cell survival.

Further strengthening this notion, we found that expression of the PhD-Polθ mutant, which cannot be phosphorylated by PLK1, failed to rescue mitotic defects occurring upon *POLQ* loss. While the expression of WT-Polθ rescued mitotic DSB accumulation, MN formation and synthetic lethality with BRCA2 loss, the expression of PhD-Polθ did not rescue these phenotypes of *POLQ^-/-^* cells (Fig. 4e-g). When challenged with RS, live microscopy revealed that *POLQ^-/-^* cells expressing PhD-Polθ showed an increased frequency of mitotic catastrophe and MN formation, together with an absence of Polθ NBs (Extended Data Fig. 9b-d and Videos 7 and 9). This indicates that PLK1-mediated regulation of mitotic Polθ repair is essential for its proper functioning.

Finally, we speculated that Polθ might be acting as part of a larger complex to repair mitotic DSBs. PARP is a known factor in alt-EJ repair that mediates Polθ recruitment to DSBs in interphase^48^. Interestingly, we found that Polθ recruitment to mitotic DSBs was abolished by PARP inhibitor (PARPi) treatment, similarly to what was reported in interphase cells^18,19^ (Extended Data Fig. 9e). One hour of PARPi treatment in mitotic *BRCA2^-/-^* cells induced the formation of DSBs, chromosomal abnormalities and was sufficient to kill *BRCA2^-/-^* but not WT cells, similarly to Polθ depletion (Extended Data Fig. 9f-h). Our data suggest that PARP, together with Polθ, enables mitotic DSB repair, a pathway crucial for HRD cell survival.

Taken together, this work explains how phosphorylation by the mitotic kinase PLK1 controls DSB repair activity in mitosis. PLK1 phosphorylation restricts the classical DSB repair pathways HR and NHEJ, but our work reveals that it enables mitotic DSB repair by Polθ. In the absence of mitotic Polθ repair, HR-deficient cells experience extreme genomic instability, leading to cell death, thus explaining the synthetic lethal relationship between Polθ and HR (Fig. 4h and Extended Fig. 9i).

## Abbreviations

Neg: negative
Pos: positive
Noc: Nocodazole
IF: immunofluorescence
Nb: number

